# Altered trajectories in the dynamical repertoire of functional network states under psilocybin

**DOI:** 10.1101/376491

**Authors:** Louis-David Lord, Paul Expert, Selen Atasoy, Leor Roseman, Kristina Rapuano, Renaud Lambiotte, David J. Nutt, Gustavo Deco, Robin L. Carhart-Harris, Morten L. Kringelbach, Joana Cabral

## Abstract

Brain activity can be understood as the exploration of a dynamical landscape of activity configurations over both space and time. This dynamical landscape may be defined in terms of spontaneous transitions within a repertoire of discrete metastable states of functional connectivity (FC), which underlie different mental processes. However, it remains unclear how the brain’s dynamical landscape might be changed in altered states of consciousness, such as the psychedelic state. The present study investigated changes in the brain’s dynamical repertoire in an fMRI dataset of healthy participants intravenously injected with the psychedelic compound psilocybin, which is found in “magic mushrooms”. We employed a data-driven approach to study brain dynamics in the psychedelic state, which focuses on the dominant FC pattern captured by the leading eigenvector of dynamic FC matrices, and enables the identification of recurrent FC patterns (“FC-states”), and their transition profiles over time. We found that a FC state closely corresponding to the fronto-parietal control system was strongly destabilized in the psychedelic state, while transitions toward a globally synchronized FC state were enhanced. These differences between brain state trajectories in normal waking consciousness and the psychedelic state suggest that the latter biases a global mode of functional integration at the expense of locally segregated activity in specific networks. These results provide a mechanistic perspective on subjective quality of the psychedelic experience, and further raise the possibility that mapping the brain’s dynamical landscape may help guide pharmacological interventions in neuropsychiatric disorders.

## Introduction

Brain function can be understood as the structured exploration of a dynamical landscape of activity configurations over both space and time (1–3). High-order mental processes involve the coordinated integration of information over spatially-distributed networks of specialized brain areas (4–8), and a robust repertoire of large-scale functional networks has been consistently detected across individuals and neuroimaging modalities (9–13). Accordingly, brain dynamics have been described in terms of spontaneous transitions within a repertoire of discrete metastable states of functional connectivity (FC), or “FC states”, which underlie different mental processes (5, 14–17).

How changes in the brain’s dynamical landscape relate to changes in behavior and cognition remains unclear however. The present study investigated changes in the brain’s dynamical repertoire in an fMRI dataset from healthy participants intravenously injected with the psychedelic compound psilocybin, all of whom reported marked changes in subjective experience during the scanning period. Investigating the link between the brain’s dynamical repertoire and departures from normal waking consciousness may help better understand functional mechanisms underlying the therapeutic potential of psilocybin for disorders including depression, anxiety and addiction (18–21). Indeed, neuropsychiatric disorders have been linked to the disruption of specific functional networks (22–25) and understanding which factors modulate the relative stability of FC states, as well as the transitions between them, may lead to the design of novel and more efficient treatments (23, 26).

It can be hypothesized that alterations in the dynamical exploration of FC states may underlie the profound departures from normal waking consciousness seen in certain altered states of consciousness. A promising experimental framework for investigating such changes is to measure brain activity in the psychedelic state. Both synthetic (i.e. LSD) and naturally-occurring psychedelic compounds (i.e. psilocybin, DMT) can profoundly alter one’s perception of their internal and external environments, and neuroimaging studies have begun to uncover the neural correlates of the psychedelic experience (27–30). Psilocybin, the compound of interest in the present study, is an ingredient in so-called ‘magic mushrooms’ and its characteristic psychoactive properties are largely mediated by activation of the serotonin 2A receptor subtype (5-HT2A receptors) (31). The phenomenology of the psilocybin experience includes a sense of ‘unconstrained’ consciousness characterized by hyper-association and profound alterations in the perception of time, space and selfhood (20, 32).

In light of these subjective effects of psilocybin, one would expect the brain’s dynamical landscape to become altered under the influence of the drug. This may be characterized by a combination of: 1) the emergence of atypical FC states, comparatively unexplored during normal wakefulness; 2) the suppression of common FC states that are normally highly visited at baseline; and/or 3) changes in the transition patterns between FC states. Preliminary support for these ideas first appeared in a fMRI study which found increased metastability in several canonical resting-state networks under psilocybin (32). A dynamical FC analysis of the same fMRI data using windowed correlations revealed both a larger repertoire of FC motifs within a small subnetwork of four brain regions comprising the bilateral hippocampus and ACC under psilocybin compared to placebo as well as greater entropy of the motif sequence (33). Moreover, a MEG study revealed increased neural signal diversity (Lempel-Ziv complexity) following psilocybin (and other psychedelic) administration relative to placebo, consistent with increased metastability (28). Finally, a recent fMRI study based on a graph signal analysis investigated the dynamical changes in brain states in healthy participants under the influence of the synthetic psychedelic LSD. It was found that the drug, which is pharmacodynamically similar to psilocybin, leads to an expansion of the repertoire of active brain states in healthy participants (34). These various findings are broadly consistent with the so-called ‘entropic brain’ hypothesis, which states that the quality of a conscious state can be indexed by the variability or randomness of its activity (32, 35).

These advances call for a detailed investigation into the dynamics of distributed functional networks in the psychedelic state that relate to its subjective properties. An improved understanding of the neural correlates underlying the experience of unconstrained cognition and enhanced mind-wandering reported by many subjects under the influence of psilocybin may provide insights into the drug’s action at the systems level and perhaps inform on its therapeutic action for the treatment of neuropsychiatric disorders such as depression (18, 30). Toward this aim, the present study employed a novel data-driven approach to study brain dynamics in the psychedelic state, which focuses on the dominant FC pattern captured by the leading eigenvector of dynamic FC matrices, and enables the identification of recurrent FC patterns over time (“FC-states”) (3). This approach enabled us to investigate the probability of occurrence of and transition profiles between FC-states extracted from fMRI data in healthy participants under the influence of psilocybin.

## Methods

### Participants

Nine healthy participants were included in this study. Study inclusion criteria were: at least 21 years of age, no personal or immediate family history of a major psychiatric disorder, substance dependence, cardiovascular disease, and no history of a significant adverse response to a psychedelic drug. All of the subjects had used psilocybin at least once before, but not within 6 weeks of the study (30). All participants gave informed consent to participate in the study. The study was approved by a National Health Service research ethics committee.

### Experimental protocol

All subjects underwent two 12 min eyes-closed resting state fMRI scans over separate sessions, at least 7 days apart. In each session, subjects were injected intravenously with either psilocybin (2 mg dissolved in 10 ml saline, 60s intravenous injection) or a placebo (10 ml saline, 60s intravenous injection) in a counterbalanced design. Injections were given manually by a doctor within the scanning suite. The infusions began exactly 6 min after the start of the 12 min scans and lasted 60 seconds. The subjective effects of psilocybin were felt almost immediately after injection and sustained for the remainder of the scanning session. This experimental approach thus provided four distinct fMRI recordings (6-min each) corresponding to each of four experimental conditions: pre/post placebo injection and pre/post psilocybin injection.

### Neuroimaging data acquisition & processing

#### Anatomical scan acquisition

Neuroimaging data were acquired using a 3T GE HDx system. Anatomical scans were performed before each functional scan and thus prior to administering either the drug or placebo. Structural scans were collected using a 3D fast spoiled gradient echo scans in an axial orientation, with field of view = 256×256×192 and matrix = 256×256×192 to yield 1 mm isotropic voxel resolution (repetition time/echo time TR/TE = 7.9/3.0 ms; inversion time = 450 ms; flip angle = 20).

#### fMRI acquisition

BOLD-weighted fMRI data were acquired using a gradient echo planar imaging sequence, TR/TE 3000/35 ms, field-of-view = 192 mm, 64 × 64 acquisition matrix, parallel acceleration factor = 2, 90 flip angle. Fifty-three oblique axial slices were acquired in an interleaved fashion, each 3 mm thick with zero slice gap (3 × 3 × 3 mm voxels). A total of 240 brain volumes were acquired, with drug/placebo infusion taking place in the middle of the session.

#### fMRI processing

fMRI data were processed data using MELODIC (Multivariate Exploratory Linear Decomposition into Independent Components) (36), part of FSL (FMRIB’s Software Library, www.fmrib.ox.ac.uk/fsl). The default parameters of this imaging pre-processing pipeline were used on all participants: motion correction using MCFLIRT (37), non-brain removal using BET (38), spatial smoothing using a Gaussian kernel of FWHM 5mm; grand-mean intensity normalization of the entire 4D dataset by a single multiplicative factor and linear de-trending over 50 second intervals. We then used tools from FSL to extract and average the time courses from all voxels within each cluster in the AAL-90 atlas (39). This procedure was applied to fMRI data from all four experimental conditions listed above and four BOLD timecourses comprising 100 TR’s each were extracted as such.

#### Dynamic Functional Connectivity Analysis

For each subject, and in each of the four experimental conditions mentioned above, we computed a phase coherence matrix (90 x 90) to capture the amount of interregional BOLD signal synchrony at each time point (TR). To compute the phase coherence between each pair of AAL regions, first the BOLD phases, θ(n,t), were estimated using the Hilbert transform for each BOLD regional timecourse. The Hilbert transform expresses a given signal x in polar coordinates as x(t)=A(t)*cos(θ(t)). Upon calculating the phases (angles) of the BOLD signals over time, the phase coherence dFC(n,p,t) between brain areas *n* and *p* at time *t* is estimated using Equation (1):

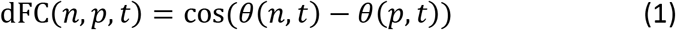

Using the cosine function, two areas n and p with temporarily aligned BOLD signals (i.e. with similar angles) at a given TR will have a phase coherence value dFC(n,p,t) close to 1 (since cos(0°)=1). On the other hand, time points where the BOLD signals are orthogonal (for instance, one increasing at 45° and the other decreasing at 45°) will have dFC(n,p,t) close to 0 (since cos(90°) = 0). The resulting dFC for each subject in each condition is a thus three-dimensional tensor with size NxNxT, where N=90 is the number of brain areas and T=100 is the total number of time points. Note that the phase coherence matrix is undirected and, as such, for each time t, the NxN dFC(t) matrix is symmetric across the diagonal.

#### FC Leading Eigenvector

To characterize the evolution of the dFC matrix over time, we employed a method termed Leading Eigenvector Dynamics Analysis (LEiDA) focusing solely on the evolution of the dominant patterns of functional connectivity, which is the partition of the ROIs into two modules/communities and captures most of the coherence based connectivity of the system (see SI for more details) focusing solely on the evolution of the dominant patterns of functional connectivity (16). The leading eigenvector, V1(t), is a Nx1 vector that captures the dominant connectivity pattern of the dFC(t) at each time t, given in matrix format by its outer product V1·V1^T^. This approach substantially reduces the dimensionality of the data when compared to more traditional approaches considering all the values in the NxN FCt(t) connectivity matrix (15, 40, 41). For further details about LEiDA, the reader is invited to consult the work of Cabral and colleagues (3) and the Supplementary Information (pp 4-7).

### FC States

#### Identification of FC states from the leading eigenvectors of phase coherence matrices

Upon computing the leading eigenvector of the phase coherence matrix dFC(t) for each TR, the next step in our analysis was to identify recurrent FC patterns in the data (‘FC states’). A discrete number of FC patterns was detected by clustering the leading eigenvectors V_1_(t) from the pooled pre/post psilocybin fMRI data (2 conditions) including all 100 TR’s of all 9 subjects, which corresponded to a combined total of 1800 V_1_(t). The k-means clustering algorithm was run with k from 2 to 20, and significant difference between the statistics of FC pre/post psilocybin was sought. Clustering solutions output a k number of cluster centroids in the shape of Nx1 V_c_ vectors for each cluster c, which represent averaged recurrent FC states. The outer product of each cluster centroid vector V_c_V_c_^T^ is a NxN rank one matrix representing the dominant connectivity pattern and the elements of V_c_ weight the contribution of each brain area to the community structure established by the signs of the corresponding vector elements. To facilitate visualization and interpretation of FC-states, the cluster centroid vectors V_c_ were rendered onto a cortical surface using HCP Workbench. For the analysis of FC states in the placebo condition, we ran the k-means clustering using the same cluster centroids obtained from the pre- and post-psilocybin injection.

#### Probability of Occurrence and Switching Matrix of FC States

Upon identifying FC states, we computed the probability of occurrence of each FC state in each condition (i.e. pre/post psilocybin and pre/post placebo). The probability of occurrence (or fractional occupancy) is simply the ratio of the number of epochs assigned to a given cluster centroid Vc divided by the total number of epochs (TRs) in each experimental condition (which is the same in all four experimental conditions). The probabilities were calculated for each subject, in each experimental condition and for the whole range of clustering solutions explored. In addition, we computed the switching matrix, which captures the trajectories of FC dynamics in a directional manner. In more detail, it indicates the probability of, being in a given FC state, transitioning to any of the other FC states.

Differences in probabilities of occurrence and probabilities of transition for the different states were statistically assessed between each pair of experimental conditions using a permutation-based paired t-test. This non-parametric test uses permutations of group labels to estimate the null distribution. The null distribution is computed independently for each experimental condition. For each of 1000 permutations a t-test is applied to compare populations and the significance threshold of α = 0.05/K was used, in order to take into account the K number of states compared in each solution. In a follow-up analysis, we also compared the number of occurrences of FC State III and FC State I at the individual subject level in the pre vs post-psilocybin injection conditions using a paired t-test.

#### Global Order and Metastability

In addition to the properties of FC states outlined above, we investigated how psilocybin injection induced alterations in the order and stability of BOLD signals. At each instant of time, the degree of order between BOLD phases *θ*(*n,t*), n=1,..,90, can be quantified using the Kuramoto Order Parameter, *OP(t)*, which can range between 0 (when all BOLD signals are out of phase) and 1 (when all BOLD signals are in phase):

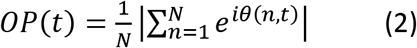

The variability of the Order Parameter is typically used to infer the degree of meta-stability in a system, whereas its mean informs whether the system is incoherent, partially synchronized or fully synchronized (42, 43). In other words, if the order parameter is constant over time, it means that the system is in a stable equilibrium, be it synchronized or not. When several attractors exist, which in this paper corresponds to the FC states, and the system switches from one to another, then the variance of the order parameter will grow with the metastability of the dynamics. To investigate the effects of psilocybin on the order and stability of BOLD activity, we therefore calculated the mean and the standard deviation of the order parameter over time, in each subject and experimental condition. Between-condition differences on these measures were assessed using a one-sample t-test.

## Results

### Dynamical repertoire of FC states is altered by psilocybin

We found a highly consistent FC pattern comprising bilateral frontal and parietal areas that had a significantly reduced probability of occurrence following the psilocybin injection, with p<0.05/k for clustering solutions in the k = 6 to k = 10 range (see Supplementary Figure S1). Over the aforementioned range of clustering solutions, this is the only FC state that reaches statistical significance between conditions upon applying the Bonferroni correction for multiple comparisons. The strongest statistical effect was observed for k = 7. The solution counting seven FC states and their associated probabilities of occurrence is plotted in Figure 1. As can be seen in the bar plots of Figure 1, the fractional occupancy of FC state III (red) significantly decreased from 14.3 ± 2.4% pre-injection (third most visited state) to 4.1 ± 1.2% following the psilocybin administration (p = 0.0003 uncorrected, p = 0.0021 after Bonferroni correction for the number of FC states compared). Conversely, a significant increase in the probability of occurrence of FC state I (dark blue) post-psilocybin injection was also observed with a fractional occupancy of 26.7 ± 4.5% pre-psilocybin injection and 37.2 ± 6.0% post-psilocybin injection (p = 0. 0068 uncorrected, p = 0.047 after Bonferroni correction for multiple comparisons). In contrast, the probability of occurrence of all other FC states remained statistically similar before and after the psilocybin injection (all p-values > 0.05). This particular FC state corresponds to a highly integrated FC state as it is the only state not presenting a natural bipartition, and is the most prevalent state across all experimental conditions (in agreement with the findings in (3). Across k = 6 to k = 10 range of clustering solutions, the fractional occupancy of this globally coherent FC state was significantly higher post-psilocybin injection and survived the Bonferroni correction for multiple comparisons for k = 7 and k = 10 (Supplementary Figure S2).

**Figure 1.**
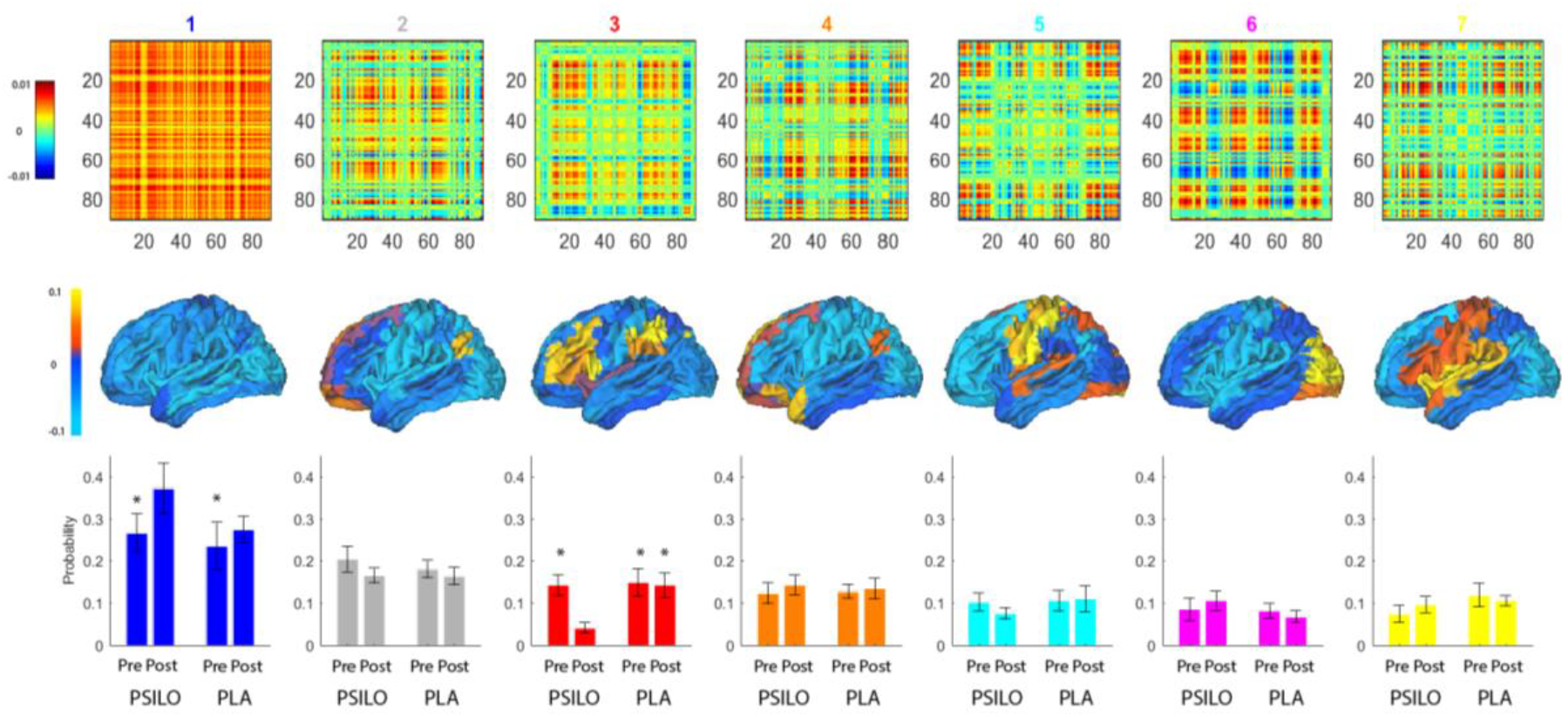
Psilocybin alters the dynamical repertoire of FC states. *Top*: Seven recurrent FC states obtained from unsupervised clustering of the eigenvectors of the dynamic FC_t_ matrix, sorted (left to right) according to decreasing probability of occurrence pre-injection. Each *NxN* matrix (given by the outer product of each cluster centroid vector *V_c_V_c_^T^*) represents a recurrent dominant connectivity pattern, *Middle:* Cluster centroid vectors *V_c_* rendered onto a cortical surface, where the N elements of *V_c_* are used to scale the color of each brain area to illustrate the contribution of each brain area to the community structure established by the signs of the corresponding vector elements. *Bottom:* Fractional occupancy of each FC state in each of the four experimental conditions (pre/post psilocybin and pre/post placebo injection). Although all FC states showed consistent probabilities of occurrence across scanning sessions in the normal resting state (i.e. pre psilocybin and pre/post placebo), we found that following the psilocybin administration the fractional occupancy of FC state III is significantly decreased from 14.3% pre-injection to 4.1% (p = 0.0077; Bonferroni corrected). Conversely, the probability of occurrence of FC state I increased after psilocybin injection (p = 0.047; Bonferroni corrected), which was not found post-placebo. The remaining FC states did not show a significant change in fractional occupancy after the psilocybin infusion in this analysis. Asterisks (*) denote the presence of a statistically significant difference (α = 0.05) in the probability of occurrence between the post-psilocybin condition and other experimental conditions for a given FC state, upon correcting for multiple comparisons.

Comparing with the separate fMRI session when a placebo solution was injected, we found that the probability of occurrence of all 7 FC states both before and after placebo injection did not differ significantly from the pre-psilocybin condition (all p-values > 0.05), indicating that the dynamical exploration of FC states assessed with LEiDA remained stable across subjects under normal conditions over the course of a week. These FC states show spatial overlap with well-known resting-state networks, namely the Default Mode Network (FC states II and IV), the Fronto-Parietal network (FC state III), the somatosensory network (FC state V), the visual network (FC state VI) and the auditory network (FC state VII). Corroborating our previous findings, the fractional occupancy of FC State III remained within normal levels both before (15.0 ± 3.3%) and after (14.3 ± 2.9%) placebo injection (p = 0.49; uncorrected), and was in each case significantly higher than the probability found during the psychedelic state (4.1 ± 1.2%, p = 0. 004 and p = 0.001 respectively, Bonferroni corrected). Moreover, the probability of FC state I, which was significantly increased after the psilocybin injection as stated above, remained unchanged before (23.6 ± 5.6%) and after the placebo injection (27.4 ± 3.1%) (p = 0.38; uncorrected). When compared to the post-psilocybin fractional occupancy of FC State I (37.2 ± 6.0%), the aforementioned pre-placebo value was significantly lower (p = 0.006; Bonferroni corrected) while the post-placebo comparison was borderline significant after correction for multiple comparisons (p = 0.07; Bonferroni corrected).

We note that the FC state whose probability of occurrence was significantly reduced following the psilocybin injection closely overlaps with the fronto-parietal network involved in executive control originally reported by Vincent and colleagues (13), which includes many regions identified as supporting cognitive control and decision-making processes such as the lateral prefrontal cortex and inferior parietal lobule (Fig. 2).

**Figure 2.**
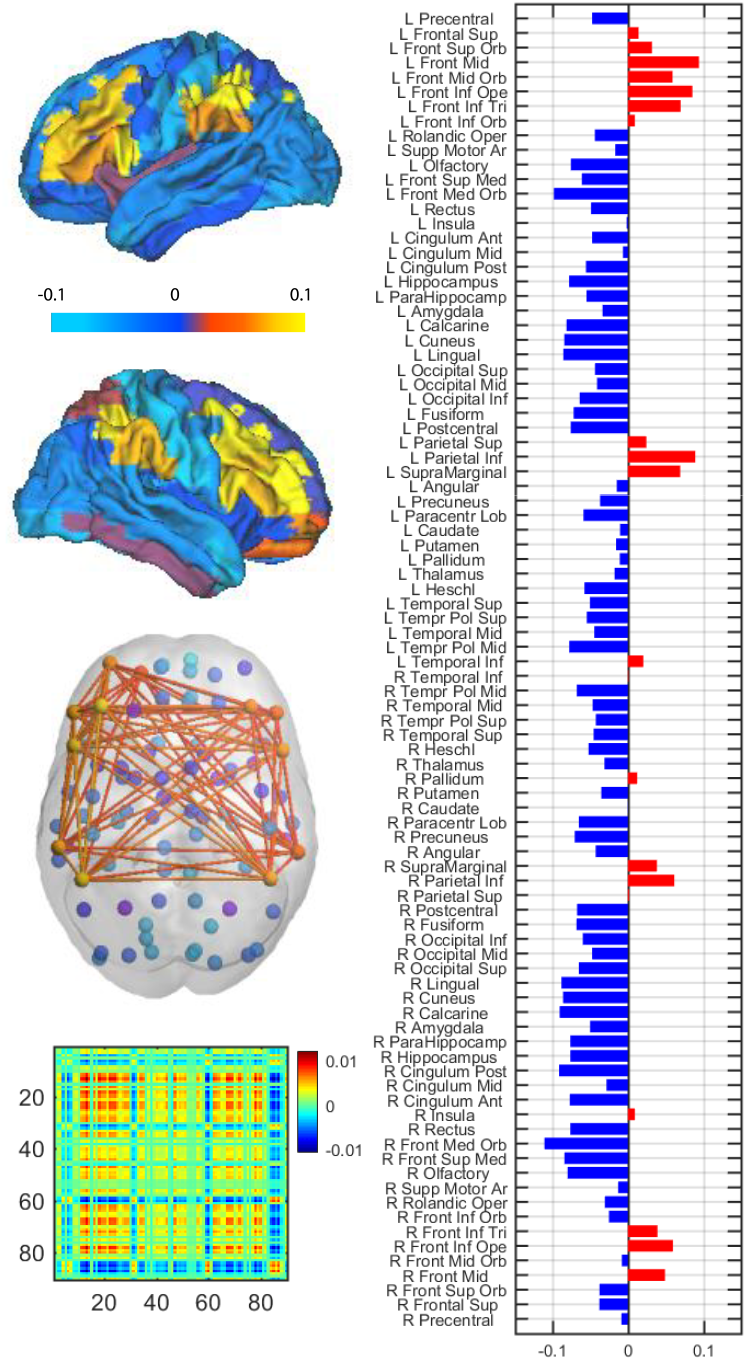
Characterization of the FC state destabilized by psilocybin. The FC state for which fractional occupancy Is significantly reduced following the psilocybin infusion corresponds to a network involving mainly bilateral frontal and parietal brain areas. Left: (top) rendition of the relevant cluster centroid vector V_c_ onto a cortical surface, highlighting the symmetry of the network of interest across both hemispheres. (middle) graphical representation of the functional network where links represent all positive functional connections between the most relevant nodes belonging to the fronto-parietal network (Vc(n)>0.02); (bottom) FC pattern represented in matrix format as the outer product of the relevant cluster centroid vector V_c_V_c_^T^. Right: Bar chart representing contributions of each of the N=90 AAL brain areas to the FC state of interest (V_c_(n)>0 in red, blue otherwise).

The non-stationary dynamics of the leading eigenvector over time reveals a constant exploration of a repertoire of FC configurations. In order to visualize this repertoire in a 3-dimensional space, we applied a PCA reduction to all leading eigenvectors across conditions and subjects (90x36000) to their first 3 Principal Components (3x36000). Then, the leading eigenvectors V_1_(t) obtained in each of the four experimental conditions were plotted as dots in this 3D space and colored according to the label assigned by the clustering algorithm. As can be seen in Figure 3 there is little overlap between clusters (also supplementary Figure S2 to view the plot from a different perspective) and all clusters are distributed around the state of global integration (blue dots). We note that, by construction, the k-means clustering algorithm produces compact clusters (i.e. non-overlapping and well separated). However these clusters originally exist in the 90 dimensional space where the leading eigenvectors live. Retaining the first 3 principal components is, however, enough to conserve well-defined clusters, as can be seen in Fig 3.

This visual rendition of the data highlights the disruption of the fronto-parietal FC state of interest (red cluster), which is clearly prevalent the 3 non-psychedelic conditions (before psilocybin and before/after placebo injection), but nearly disappears after the psilocybin injection (upper right-hand corner). It can also be seen that the blue cluster corresponding to a globally coherent FC state becomes denser after the psilocybin injection (Fig. 3).

**Figure 3.**
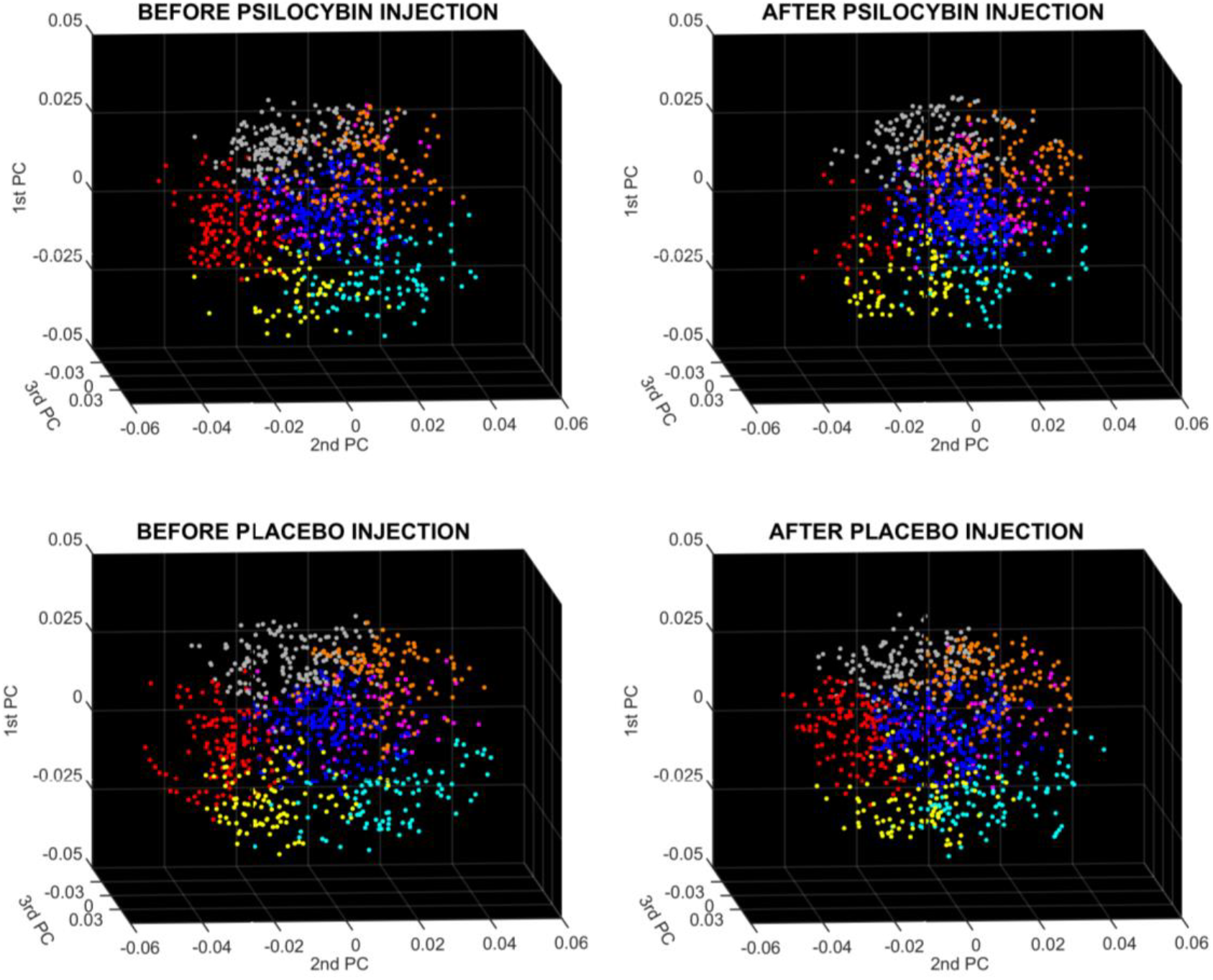
3D embedding of FC patterns reveals the reduced stability of the fronto-parietal network (red) after psilocybin injection.

In each scatter plot, each dot represents an FC pattern, *V_1_(t),* occurring during the corresponding session (before/after psilocybin/placebo, with 900 dots per subplot), projected onto the first 3 principal components of all eigenvectors across sessions (PCA-reduced from 90x3600 to 3x3600). Dots are colored according to the *k*-means clustering solution obtained before using the same color code as in Figure 1. It can be clearly observed that after the psilocybin injection the number of red dots is reduced indicating less detections of the frontal-parietal network (shown in Figure 2). Moreover, the network in the center of the 3D scatter plots corresponds to the global mode (blue), and all the remaining FC states appear as clouds of dots around it. See Supplementary Figure S3 for a different perspective of the same 3D scatter plots.

#### Psilocybin increases whole-brain metastability and modulates the temporal expression of FC States

We found a significant increase in whole-brain metastability following the psilocybin injection (0.18 ± 0.01) as reflected by an increase in the standard deviation of the order parameter of the system over time relative to the pre-injection baseline (0.16 ± 0.015) ; t(8) = 2.41, p = 0.04 (Fig. 4, bottom panel). On the other hand, no significant difference was found when considering the mean of the order parameter over time, which reflects the average level of synchrony across the nodes comprised in the whole-brain network throughout the entire recording interval. We also plotted, for each subject, the evolution of the order parameter of the system at each of the 100 time points pre/post-psilocybin injection, as well as the corresponding leading eigenvector V1(t) color-coded according to their respective cluster assignment (Fig. 4, top panel). In all subjects, the eigenvector corresponding to the global signal (1st state from top; blue colored) is the most frequently expressed both before and after the psilocybin injection, and its fractional occupancy is particularly elevated in subjects with the highest average order parameter values over time. We further found that, with the exception of subject *s1,* the fronto-parietal FC state of interest (3rd state from top; red-colored) was approximately evenly expressed in all subjects in the pre-injection phase. Following the psilocybin injection, the fractional occupancy of this state was decreased in all subjects. We note that, in the case of subjects *s4* and *s9,* not a single V1(t) in the post-psilocybin injection condition belongs to the fronto-parietal FC state. This was indeed the only FC state where a fractional occupancy of zero was found amongst all subjects and experimental conditions. A paired-sample t-test revealed that, at the individual subject level, the mean number of occurrences of FC state III (fronto-parietal network) was significantly decreased following the psilocybin injection (4.1 ± 1.2) relative to the pre-injection baseline (14.3 ± 2.5); t(8) = 3.88, p = 0.005). Conversely, a paired-sample t-test showed a significant *increase* in the mean number of occurrences of FC state I (globally integrated state) following the psilocybin injection (37.2 ± 6.0) relative to the pre-injection baseline (26.7 ± 4.5); t(8) = 2.75, p = 0.01 (Fig. 4, bottom panel).

**Figure 4.**
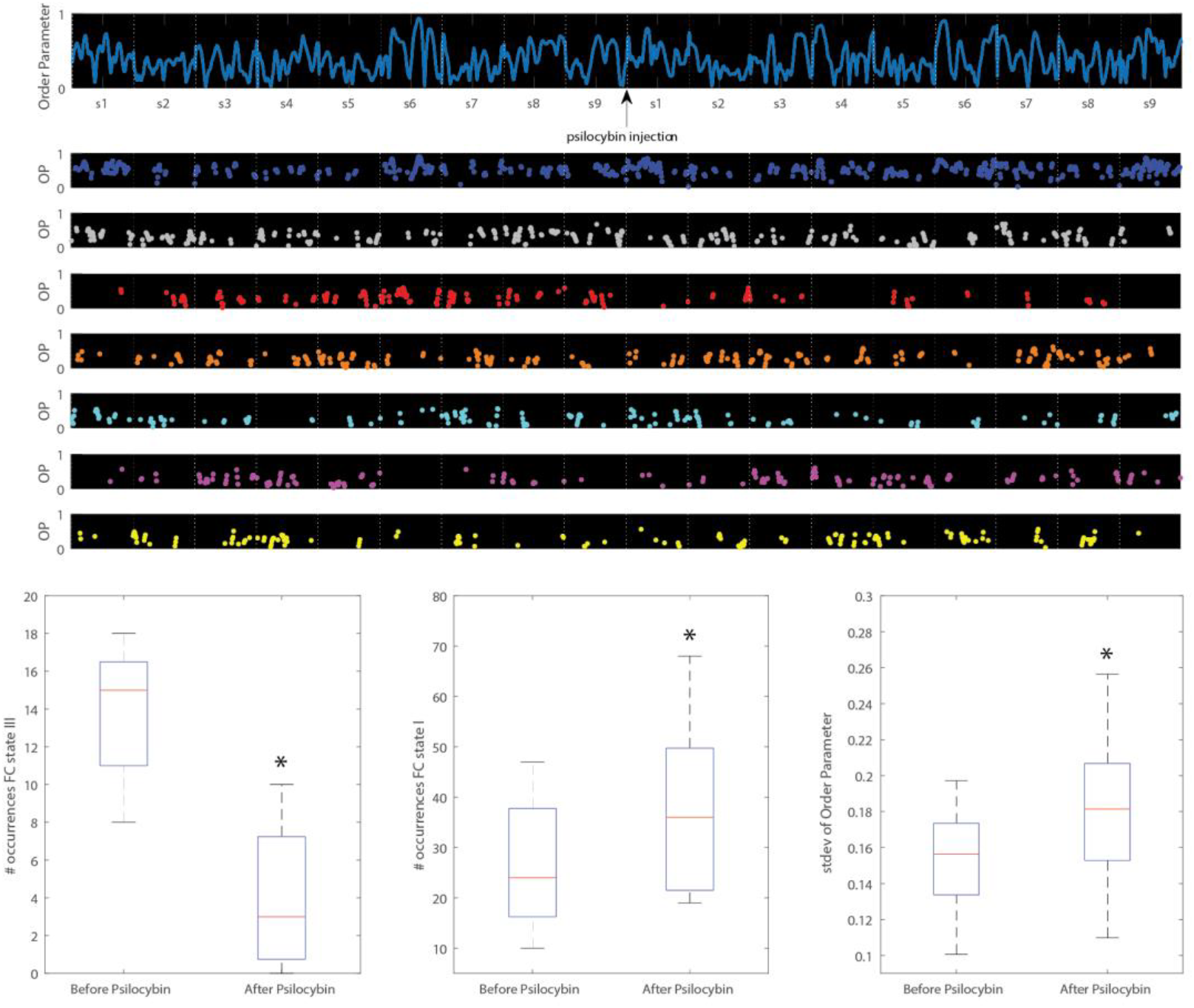
Psilocybin modulates the temporal expression of FC States and increases whole-brain metastability. *Top:* For each of the 9 subjects *(s1, s2… s9),* we show the order parameter (OP) of the system over time pre/post-psilocybin injection (100 TR/ subject/condition). Below, we show the order parameter when each of the seven FC states in ON (same color-code as Figure 1, sorted vertically by decreasing probability pre-injection). In all subjects, the FC state corresponding to the global signal (blue colored) is the most frequently expressed, and the order parameter is particularly high when this state is ON. On the other hand, less-frequent states, namely the light blue, purple, or yellow, correspond to epochs of lower order parameter values. Following the psilocybin injection, the red-colored fronto-parietal FC state of interest (3^rd^ most prevalent state pre-injection) exhibits a significant decrease in all subjects. *Bottom:* The average number of occurrences of FC State III (fronto-parietal network) across subjects is significantly lower after the psilocybin injection compared to the pre-injection baseline (left), while the average number of occurrences of FC State I (globally integrated state) is significantly increased (middle). A significant increase in whole-brain metastability following the psilocybin injection, as reflected by an increase in the standard deviation of the order parameter of the system over time (right).

#### State-to-State switching profiles are significantly different under psilocybin

The switching matrices shown in Fig. 5 (top) display the probability of: 1) being in a given FC state (rows), and 2) transitioning to any of the other states (columns) both before and after the psilocybin injection. An illustration of pre-vs post-injection changes in the transitions probabilities between FC-states rendered on the cortical surface is also provided to facilitate the neuroanatomical interpretation of FC state switching (Fig. 5, bottom). We find that the probability of transitioning from any given FC state to the fronto-parietal FC state of interest is consistently reduced post-psilocybin injection than prior to the injection. Conversely, all FC states except for one were more likely to transition toward the FC state corresponding to the global signal following the psilocybin injection. The p-values for each state-to-state transition in the switching matrix are provided in Supplementary Table ST1.

**Figure 5.**
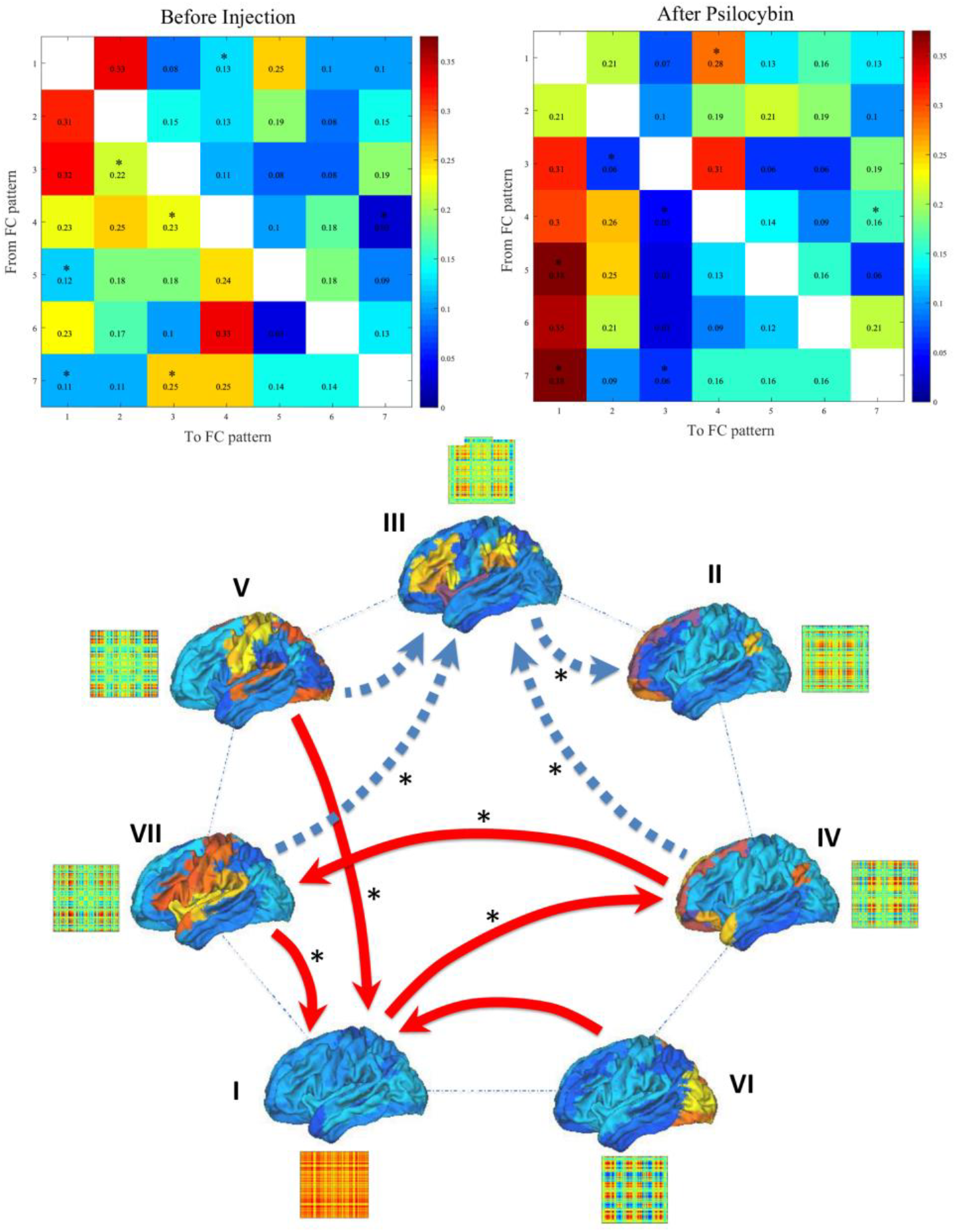
Psilocybin modifies the switching patterns between FC-states. *Top:* Switching matrices showing the probability of, being in a given FC state (rows), transitioning to any of the other states (columns) both before (left) and after the psilocybin injection (right). Significant between-condition differences assessed via a permutation test are indicated by asterisks (*) for the significance threshold α = 0.05. *Bottom:* Pre vs post-injection changes in the transitions probabilities between FC-states rendered on the cortical surface, and numbered according to the transition matrices above. Each arrow represents a state-to-state transition probability ± 1 SD from the mean change in transition probability post-injection; red/filled arrows represent a greater probability of transition post-injection, while blue/dashed arrows show reduced transition probabilities between states. It can be observed that the probability of transitioning from several FC states *(IV, V, VII)* to the frontoparietal FC state of interest is reduced post-psilocybin injection. Conversely, most FC states (*IV*, *V*, *VI*, *VII*) were more likely to transition toward the globally integrated FC state I following the psilocybin injection.

## Discussion

The present results provide the first experimental evidence that the brain’s dynamical landscape is altered by the psychedelic compound psilocybin. We found that the repertoire of functional networks at rest (described as “FC states”), and the transitions between them, are significantly changed by the drug. Notably, the strongly reduced fractional occupancy of a FC state overlapping with the previously-described fronto-parietal control system suggests that this particular functional network becomes destabilized under psilocybin (13). Conversely, transitions toward a globally integrated FC state were enhanced under the influence of the drug. The destabilization of a frontoparietal FC state supporting executive function, and increased transitions toward a highly integrated brain state provide a novel mechanistic perspective into psilocybin effects at the systems level. Indeed, these differences between brain state trajectories in normal waking consciousness and the psychedelic state suggest that the drug induces an alternative type of functional integration at the expense of locally segregated activity in specialized networks. These new insights into altered brain dynamics in the psychedelic state may also help explain some the acute psychological effects of psilocybin, and the compound’s promising therapeutic applications.

The decreased probability of occurrence of a FC state closely overlapping with the fronto-parietal control system is consistent with prior human neuroimaging studies of psilocybin effects. An fMRI study of the same dataset notably revealed that the psilocybin infusion modified the BOLD spectral content in a distributed fronto-parietal network. Specifically, the fronto-parietal system of interest showed a significantly decreased low frequency power and power spectrum scaling exponent following the psilocybin infusion (33). Similarly, a MEG study reported decreased oscillatory power following a psilocybin injection in a bilateral fronto-parietal network derived from beta-band activity (44). Findings from these prior studies indicate a desynchronization of the fronto-parietal control system under psilocybin and are thus consistent with the present results. Furthermore, the disengagement of the frontoparietal system can be interpreted in light of the cognitive and behavioral effects of psilocybin reported by the study participants. Evidence suggests the fronto-parietal system plays a key role in cognitive control by focusing attention on goal-relevant information through top-down mechanisms (45–47). Declines in fronto-parietal FC have been associated with difficulty suppressing goal-irrelevant information, and greater distractibility (48). These observations are broadly consistent with the reports of increased mind-wandering and unconstrained cognition reported by participants in the present cohort (30).

The present study also found increased FC state transitions toward a globally synchronized functional network following the psilocybin injection. Previous work has shown that psilocybin induces an alternative type of functional integration, characterized by greater global integration (27, 49) as well as increases in the entropy or metastability of brain states (32). The results presented here not only align with these prior findings but also shed further light on their functional significance. Here we found that a globally integrated state, i.e. FC state I, increased its predominance under psilocybin, not only statistically over time (Fig. 1), but also dynamically, as was evident from the structure of the state-to-state switching matrix (Fig. 5), where it became the ‘state of choice’ for inter-state transitions. In other words, under psilocybin the brain spent more time in globally coherent, highly integrated state. This state therefore served as an attractor, absorbing the majority of inter-state transitions. As was implied by previous analyses, this effect may enable atypical patterns of interregional communication to arise in the psychedelic state (27) alongside the breakdown of within-network integrity in favor of globally integrated brain dynamics (27, 32, 44, 49, 50).

In addition to changing the relative stability of FC states and the directionality of state transitions, we found that psilocybin also increases the global metastability of brain dynamics. This result is consistent with prior fMRI investigations of the same psilocybin dataset in which a greater diversity of functional connectivity motifs was observed after psilocybin infusion within a small network consisting of the bilateral hippocampal and ACC regions, thus reflecting increased variability in the collective repertoire of metastable states (33). This prior analysis had purposely limited the number of regions included in the analysis in order to perform an exhaustive counting of all possible FC states. Another study had found increased metastability in several canonical resting-state networks as determined by the variance in the network’s intrinsic synchrony over time (relationship between the mean and variance of the signal over time) (32). Here we used a complementary analytical approach to demonstrate that increased metastability of the FC under psilocybin is generalizable at the whole-brain scale by calculating the standard deviation of the order parameter of the system over time. The increased metastability of FC dynamics over time under psilocybin may be interpreted in the context of the “entropic brain” theory of conscious states according to which more variable brain activity and dynamics reflect more a variable and dynamic conscious experience (32, 34, 35).

Despite the relatively small number of subjects (N=9) included in this fMRI dataset, our findings are strongly validated by the replication of results in placebo conditions across the two scanning days (i.e. no changes in the distribution of network states in the placebo scanning session vs pre-psilocybin infusion), as well as the strong statistical effect of decreased probability of the FC state of interest after psilocybin injection (p < 0.0005). Importantly, since LEiDA compares the statistics between whole-brain FC patterns rather than individual functional links, the dimensionality is reduced from N(N-1)/2 (number of unique links in a NxN symmetric matrix) to the number of networks K, which overcomes several of the statistical limitations associated with multiple comparisons. Furthermore, given the instantaneous nature of LEiDA, our dynamic FC analysis revealed significant between-condition differences despite the fact that each scanning condition consisted only of 100 fMRI volumes, which would likely not have been sufficient for conventional sliding-window analysis (51). We note however, that the present analysis may not capture the fast dynamics of functional networks, which are believed to have lifetimes of approximately 200ms as suggested by MEG studies (52, 53), but still captures changes in their probability of occurrence over slower time scales.

The therapeutic potential of psilocybin in psychiatry has recently generated much interest. For example, indications of efficacy have been reported for conditions including treatment-resistant depression, anxiety related to end-of-life care and addictive disorders (18–21). The neural mechanisms underlying these clinical benefits remain unclear however. The present results provide some of the first evidence that a psychoactive drug can modulate the brain’s dynamical landscape by selectively destabilizing a particular brain state, and promoting the transitions towards another, namely a globally integrated brain state. Importantly, the fact that a 5-HT2A receptor agonist can be used to target specific brain states in this way opens up the possibility of studying the effects of other classes of neuromodulators at the systems level and how they may be involved in the pathophysiology and potential treatment of neuropsychiatric disorders. In the present case, a FC state closely overlapping with the frontoparietal control system was selectively disengaged following the administration of psilocybin. This raises exciting expectations for the design of novel therapeutics for neuropsychiatric disorders informed by patho-connectomics, which may be generally understood in terms of targeting the specific FC states affected by a particular disorder.

In summary, the present study used a novel, data-driven dynamical functional connectivity analysis (LEiDA) to help uncover the dynamics between functional connectivity states under the influence of psilocybin. We found that a FC state closely corresponding to the fronto-parietal control system was strongly destabilized by the compound, while transitions toward a globally synchronized FC state were enhanced. We also found an increase in the metastability of global brain dynamics following the psilocybin infusion. Taken together, these findings are consistent with prior neuroimaging studies suggesting that a different type of brain integration and increased neural signal complexity underlie the psychedelic state. The present results also raise the possibility that mapping the brain’s dynamical landscape may help guide pharmacological interventions in neuropsychiatric disorders by targeting the dynamical trajectories of functional brain states.

## Acknowledgments

LDL is supported by the Canadian Institutes of Health Research, Canadian Centennial Scholarship Fund and the Mann Senior Scholarship from Hertford College, University of Oxford. MLK is supported by the ERC Consolidator Grant CAREGIVING (n. 615539) and Center for Music in the Brain, funded by the Danish National Research Foundation (DNRF117). JC is supported under the project NORTE-01-0145-FEDER-000023 from the Northern Portugal Regional Operational Programme (NORTE 2020) under the Portugal 2020 Partnership Agreement through the European Regional Development Fund (FEDER). RCH is supported by the Alex Mosley Charitable Trust. The authors are grateful to the Beckley Foundation for funding the original research study, on which Amanda Feilding, Director of the Beckley Foundation was a collaborative partner.

## References

1. Gu S, Cieslak M, Baird B, Muldoon SF, Grafton ST, Pasqualetti F, et al. The energy landscape of neurophysiological activity implicit in brain network structure. Scientific reports. 2018;8(1):2507.

2. Watanabe T, Hirose S, Wada H, Imai Y, Machida T, Shirouzu I, et al. Energy landscapes of resting-state brain networks. Frontiers in neuroinformatics. 2014;8:12.

3. Cabral J, Vidaurre D, Marques P, Magalhães R, Moreira PS, Soares JM, et al. Cognitive performance in healthy older adults relates to spontaneous switching between states of functional connectivity during rest. Scientific Reports. 2017;7(1):5135.

4. Varela FJ. Principles of biological autonomy. 1979.

5. Cavanna F, Vilas MG, Palmucci M, Tagliazucchi E. Dynamic functional connectivity and brain metastability during altered states of consciousness. NeuroImage. 2017.

6. Menon V. Large-scale brain networks and psychopathology: a unifying triple network model. Trends in cognitive sciences. 2011;15(10):483–506.

7. Bressler SL, Menon V. Large-scale brain networks in cognition: emerging methods and principles. Trends in cognitive sciences. 2010;14(6):277–90.

8. Lord L-D, Stevner AB, Deco G, Kringelbach ML. Understanding principles of integration and segregation using whole-brain computational connectomics: implications for neuropsychiatric disorders. Philosophical Transactions of the Royal Society A: Mathematical, Physical and Engineering Sciences. 2017;375(2096).

9. Seeley WW, Menon V, Schatzberg AF, Keller J, Glover GH, Kenna H, et al. Dissociable intrinsic connectivity networks for salience processing and executive control. The Journal of neuroscience : the official journal of the Society for Neuroscience. 2007;27(9):2349–56.

10. Brookes MJ, Hale JR, Zumer JM, Stevenson CM, Francis ST, Barnes GR, et al. Measuring functional connectivity using MEG: methodology and comparison with fcMRI. NeuroImage. 2011;56(3):1082–104.

11. Hipp JF, Hawellek DJ, Corbetta M, Siegel M, Engel AK. Large-scale cortical correlation structure of spontaneous oscillatory activity. Nature neuroscience. 2012;15(6):884–90.

12. Menon V, Uddin LQ. Saliency, switching, attention and control: a network model of insula function. Brain structure & function. 2010;214(5–6):655–67.

13. Vincent JL, Kahn I, Snyder AZ, Raichle ME, Buckner RL. Evidence for a frontoparietal control system revealed by intrinsic functional connectivity. Journal of neurophysiology. 2008;100(6):3328–42.

14. Tononi G, Edelman GM. Consciousness and complexity. science. 1998;282(5395):1846–51.

15. Hansen EC, Battaglia D, Spiegler A, Deco G, Jirsa VK. Functional connectivity dynamics: modeling the switching behavior of the resting state. NeuroImage. 2015;105:525–35.

16. Cabral J, Vidaurre D, Marques P, Magalhaes R, Moreira P, Soares J, et al. Cognitive performance in healthy older adults relates to spontaneous switching between states of functional connectivity during rest. Scientific reports. 2017;In press.

17. Cabral J, Kringelbach M, Deco G. Functional connectivity dynamically evolves on multiple time-scales over a static structural connectome: Models and mechanisms. NeuroImage. 2017.

18. Carhart-Harris RL, Bolstridge M, Rucker J, Day CM, Erritzoe D, Kaelen M, et al. Psilocybin with psychological support for treatment-resistant depression: an open-label feasibility study. The Lancet Psychiatry. 2016;3(7):619–27.

19. Johnson MW, Garcia-Romeu A, Cosimano MP, Griffiths RR. Pilot study of the 5-HT2AR agonist psilocybin in the treatment of tobacco addiction. Journal of psychopharmacology. 2014;28(11):983–92.

20. Griffiths RR, Richards WA, McCann U, Jesse R. Psilocybin can occasion mystical-type experiences having substantial and sustained personal meaning and spiritual significance. Psychopharmacology. 2006;187(3):268–83.

21. Grob CS, Danforth AL, Chopra GS, Hagerty M, McKay CR, Halberstadt AL, et al. Pilot study of psilocybin treatment for anxiety in patients with advanced-stage cancer. Archives of general psychiatry. 2011;68(1):71–8.

22. Fornito A, Zalesky A, Breakspear M. The connectomics of brain disorders. Nature reviews Neuroscience. 2015;16(3):159–72.

23. Deco G, Kringelbach ML. Great expectations: using whole-brain computational connectomics for understanding neuropsychiatric disorders. Neuron. 2014;84(5):892–905.

24. Stam CJ. Modern network science of neurological disorders. Nature reviews Neuroscience. 2014;15(10):683–95.

25. Hellyer PJ, Scott G, Shanahan M, Sharp DJ, Leech R. Cognitive Flexibility through Metastable Neural Dynamics Is Disrupted by Damage to the Structural Connectome. The Journal of neuroscience : the official journal of the Society for Neuroscience. 2015;35(24):9050–63.

26. Kringelbach ML, Green AL, Aziz TZ. Balancing the brain: resting state networks and deep brain stimulation. Frontiers in integrative neuroscience. 2011;5:8.

27. Petri G, Expert P, Turkheimer F, Carhart-Harris R, Nutt D, Hellyer PJ, et al. Homological scaffolds of brain functional networks. Journal of The Royal Society Interface. 2014;11(101):20140873.

28. Schartner MM, Carhart-Harris RL, Barrett AB, Seth AK, Muthukumaraswamy SD. Increased spontaneous MEG signal diversity for psychoactive doses of ketamine, LSD and psilocybin. Scientific Reports. 2017;7.

29. Carhart-Harris RL, Muthukumaraswamy S, Roseman L, Kaelen M, Droog W, Murphy K, et al. Neural correlates of the LSD experience revealed by multimodal neuroimaging. Proceedings of the National Academy of Sciences. 2016;113(17):4853–8.

30. Carhart-Harris RL, Erritzoe D, Williams T, Stone JM, Reed LJ, Colasanti A, et al. Neural correlates of the psychedelic state as determined by fMRI studies with psilocybin. Proceedings of the National Academy of Sciences. 2012;109(6):2138–43.

31. González-Maeso J, Weisstaub NV, Zhou M, Chan P, Ivic L, Ang R, et al. Hallucinogens recruit specific cortical 5-HT 2A receptor-mediated signaling pathways to affect behavior. Neuron. 2007;53(3):439–52.

32. Carhart-Harris RL, Leech R, Hellyer PJ, Shanahan M, Feilding A, Tagliazucchi E, et al. The entropic brain: a theory of conscious states informed by neuroimaging research with psychedelic drugs. Frontiers in human neuroscience. 2014;8.

33. Tagliazucchi E, Carhart-Harris R, Leech R, Nutt D, Chialvo DR. Enhanced repertoire of brain dynamical states during the psychedelic experience. Human brain mapping. 2014;35(11):5442–56.

34. Atasoy S, Roseman L, Kaelen M, Kringelbach ML, Deco G, Carhart-Harris RL. Connectome-harmonic decomposition of human brain activity reveals dynamical repertoire re-organization under LSD. Scientific reports. 2017;7(1):17661.

35. Carhart-Harris RL. The entropic brain-Revisited. Neuropharmacology. 2018.

36. Beckmann CF, Smith SM. Probabilistic independent component analysis for functional magnetic resonance imaging. IEEE transactions on medical imaging. 2004;23(2):137–52.

37. Jenkinson M, Bannister P, Brady M, Smith S. Improved optimization for the robust and accurate linear registration and motion correction of brain images. Neuroimage. 2002;17(2):825–41.

38. Smith SM. Fast robust automated brain extraction. Human brain mapping. 2002;17(3):143–55.

39. Tzourio-Mazoyer N, Landeau B, Papathanassiou D, Crivello F, Etard O, Delcroix N, et al. Automated anatomical labeling of activations in SPM using a macroscopic anatomical parcellation of the MNI MRI single-subject brain. Neuroimage. 2002;15(1):273–89.

40. Hutchison RM, Womelsdorf T, Allen EA, Bandettini PA, Calhoun VD, Corbetta M, et al. Dynamic functional connectivity: promise, issues, and interpretations. NeuroImage. 2013;80:360–78.

41. Preti MG, Bolton TA, Ville DV. The dynamic functional connectome: State-of-the-art and perspectives. NeuroImage. 2016.

42. Deco G, Kringelbach M. Metastability and Coherence: Extending the Communication through Coherence Hypothesis Using a Whole-Brain Computational Perspective. Trends in neurosciences. 2016;39(6):432.

43. Cabral J, Hugues E, Sporns O, Deco G. Role of local network oscillations in resting-state functional connectivity. Neuroimage. 2011;57(1):130–9.

44. Muthukumaraswamy SD, Carhart-Harris RL, Moran RJ, Brookes MJ, Williams TM, Errtizoe D, et al. Broadband cortical desynchronization underlies the human psychedelic state. Journal of Neuroscience. 2013;33(38):15171–83.

45. Xin F, Lei X. Competition between frontoparietal control and default networks supports social working memory and empathy. Social cognitive and affective neuroscience. 2015;10(8):1144–52.

46. Cole MW, Schneider W. The cognitive control network: integrated cortical regions with dissociable functions. Neuroimage. 2007;37(1):343–60.

47. Dosenbach NU, Fair DA, Cohen AL, Schlaggar BL, Petersen SE. A dual-networks architecture of top-down control. Trends in cognitive sciences. 2008;12(3):99–105.

48. Smallwood J, Brown K, Baird B, Schooler JW. Cooperation between the default mode network and the frontal-parietal network in the production of an internal train of thought. Brain research. 2012;1428:60–70.

49. Roseman L, Leech R, Feilding A, Nutt DJ, Carhart-Harris RL. The effects of psilocybin and MDMA on between-network resting state functional connectivity in healthy volunteers. Frontiers in human neuroscience. 2014;8:204.

50. Tagliazucchi E, Roseman L, Kaelen M, Orban C, Muthukumaraswamy SD, Murphy K, et al. Increased global functional connectivity correlates with LSD-induced ego dissolution. Current Biology. 2016;26(8):1043–50.

51. Hindriks R, Adhikari MH, Murayama Y, Ganzetti M, Mantini D, Logothetis NK, et al. Can sliding-window correlations reveal dynamic functional connectivity in resting-state fMRI? NeuroImage. 2015;127:242–56.

52. Baker AP, Brookes MJ, Rezek IA, Smith SM, Behrens T, Smith PJP, et al. Fast transient networks in spontaneous human brain activity. eLife. 2014;3.

53. Vidaurre D, Quinn AJ, Baker AP, Dupret D, Tejero-Cantero A, Woolrich MW. Spectrally resolved fast transient brain states in electrophysiological data. NeuroImage. 2016;126:81–95.

